# Computational Assessment of Aromatase: Unmasking Therapeutic Avenues for Female Hormonal Disorders

**DOI:** 10.1101/2023.09.13.557603

**Authors:** Sandesh P. Koirala, Bhawana S. Magar, Amit K. Yadav, Anisha Regmi, Aashish Pokharel, Samiran Subedi, Pramod Aryal

## Abstract

Women of reproductive age widely experience the physical, mental, and emotional effects of hormonal imbalances. The levels of two hormones, estrogen, and progesterone, were particularly high. When these two hormones are unbalanced, symptoms such as irregular menstrual cycles, pelvic pain, and uterine fibroids can appear. Statistics show that 80 percent of women have hormonal imbalances. The creation of novel disease-containing drug candidates may be facilitated by computer-aided drug design (CADD), as many drugs and surgical treatments have unique adverse effects. Achieving good data coverage requires an optimized process for protein identification and enhanced selection techniques for medicinal compounds, including their metabolic characterization. In this study, aromatase is the prioritized protein that acted as a rate-limiting enzyme in steroidogenesis. To maintain its expression and regulation to control estrogen production, natural compounds were retrieved from the ZINC15 database through ADME/TOX property filtration. Compound B, specifically designated as (1R,3S,6S,7S,10S,11R)-7-[(dimethylamino)methyl]-3,12-dimethyl-2,9-dioxatetracyclo [9.3.0.01,3.06,10] tetradec-12-en-8-one, exhibits remarkable therapeutic potential, as evidenced by its inability to inhibit S-adenosyl methionine (SAM) biosynthesis, the observed stability of molecular interactions based on Density Functional Theory (DFT), and a favorable energy gap between molecular orbital stages. This research can be employed as a utility module for fast screening of drug-like molecules and taking these leads forward to clinical trials, assessing their value as potential suitors for drug repurposing against female hormonal disorders.

## Introduction

WHO defines “health” as a state of complete physical, mental, and social well-being and not merely the absence of disease or infirmity. (World Health Organization [WHO], 1946). Menstrual health is an essential part of women’s overall health, as it has a significant impact on their physical, mental, and social well-being (Matteson et al., 2013). Any cyclic blood loss that occurs from the uterine corpus that take place between menarche and menopause with the average is menstruation. In cycles between 21 and 35 days, the average menstrual blood loss is 7 days with the average amount of blood lost during a period is 30 to 40 ml, with the first few days of the period showing the greatest blood loss is called normal menstruation (Munro et al., 2018).

Menstrual bleeding is not always “normal” for all women. (Kjerulff et al., 1996). According to Shah (2017), a female encounters an array of menstrual disorders from puberty to menopause. Early adolescent and young adult females showed a prevalence and pattern of menstrual disorders. Up to 30% of women experience variations in the volume or pattern of menstrual blood flow, referred to as the symptom of abnormal uterine bleeding (AUB), which can be brought on by a number of etiologies, sometimes more than one (Kjerulff et al., 1996). Heavy and irregular menstrual bleeding are the two main subtypes of AUB, and many patients report a combination of these symptoms (Lobo et al., 2016).

Any blood loss of ≥ ml or more and/or a duration of > 7 days was classified as heavy menstrual bleeding (HMB) or menorrhagia (Tidy, 2020), (Rees, 1987). Menorrhagia is the most common menstrual disorder among women (Philipp et al., 2005). There are various potential causes of HMB, including ovulatory abnormalities, adenomyosis, endometriosis, endometrial polyps, and endometrial hyperplasia; nevertheless, uterine fibroids are the most frequently occurring conditions underlying HMB (Stewart et al., 2016).

Uterine fibroids, or leiomyomas, are the most common benign tumors of the myometrium that arise within the uterus (Holdsworth-Carson et al., 2016). More than 70% of women globally, especially women of color, are affected by uterine fibroids, the most prevalent pelvic tumor among women of reproductive age (Baird et al., 2003) (Wise & Laughlin-Tommaso, 2016) (Al-Hendy et al., 2017).

Uterine fibroids are benign tumors that grow throughout the reproductive years and are often controlled during menopause. They are sex steroid hormone-dependent. In addition to the ovary, the leiomyomal tissue itself provides an oestrogen source. Leiomyomal tissue oestrogen secretion may accumulate to a sufficiently high level in the local compartment to support its own growth, enabling independence from ovarian oestrogen. Additionally, uterine fibroids have higher levels of aromatase and 17b-hydroxysteroid dehydrogenase type I than the myometrium (Bulun et al., 2005).

Circulating androstenedione is transformed into estrone and subsequently into estradiol, the active form of estrogen, by the enzyme 17b-HSD type I in leiomyoma cells (Sumitani et al., 2000). In this process, aromatase is a rate-limiting enzyme in the conversion of C19 androgenic steroids to the corresponding estrogens hence, it appears to be an important enzyme (Bulun et al., 1994) (Kamat et al., 2002).

Traditionally, uterine fibroids are treated surgically. Surgery can be avoided by increasing the demand for medical treatment, protecting the uterus, and improving potential future fertility. For the majority of women, a comfortable and non-invasive method of reducing fibroids could be desirable, and it might also ease the strain on the healthcare systems. AIs have been the subject of several clinical investigations for the treatment of uterine fibroids (Gurates et al., 2008) (Hil á rio et al., 2009).

Aromatase inhibitors effectively block the in-situ production of estradiol, thus decreasing leiomyoma or uterine fibroids’ responsiveness to both estrogen and progesterone signaling (Bulun et al., 1994). Drug aromatase inhibitors do not prevent estrogen production in ovaries. Instead, they only lower estrogen levels in women whose ovaries do not contain estrogen. Therefore, only women who have experienced menopause are eligible to use such drugs (Breast Cancer Prevention: Aromatase Inhibitors, n.d.).

Disturbances in regular menstrual cycles may present diagnostic and management challenges for gynecologists and contribute to future disturbances for adolescents and families (Patil et al., 2017). The National Institute for Health and Care Excellence (NICE) defines heavy menstrual loss as excessive blood loss that interferes with a woman’s physical, social, emotional, and/or quality of life (NICE, 2018). Therefore, to address the drug-ability and prophylaxis of menorrhagia in women of reproductive age, multi-target proteins need to be prioritized, which can be achieved by protein-protein interactions (PPI). PPI, on the other hand, generates multiple targets, making it suitable for identifying combinational therapies.

Despite numerous advances in the ability to prevent or postpone cell damage, the anticipated number of promising New Medical Entities (NME) needed to control disease progression and completely eradicate it has not yet been attained. To effectively build a new methodology to find targets, it may be possible to identify potential targets and their role in disease-related signaling using functional protein-protein interactions. For effective treatment with few or no adverse effects, Virtual High-throughput Screening (VHTS) of a novel drug candidate is required, preferably derived from natural materials, using Computer Aided Drug Discovery (CADD).

This work is solely based on the application of several techniques concerning CADD for the development of potential drug candidates maintaining the aromatase enzyme at its normal threshold.

## Results

### Protein Structural Retrieval, Validation, and Active Site Generation

Aromatase is a rate-limiting step in the hormonal pathway for estrogen production (Jeong et al., 2010). The crystal structure of the aromatase protein (PDB ID:3EQM) catalyzes the biosynthesis of all estrogens from androgens (Ghosh et al., 2009). It consists of a hem group and a polypeptide chain of 503 amino acid residues and has a molecular weight of 57.95kDa. It has an ‘A’ chain with a sequence length of 503 (Ghosh et al., 2009). A good protein model usually has 90% of its residues in the allowable regions of the Ramachandran plot (Laskowski et al., 1993). However, the target protein structure that was deposited comprised 86.6% of the residues in the allowed regions. Because of the proximity of the target protein to 90%, it was considered a good model of the protein.

For aromatase, the active binding sites were determined by taking the amino acid residues lying within 5 A of the protein and native ligand interaction sites which can be observed in *Figure 1*.

**Figure 1:**
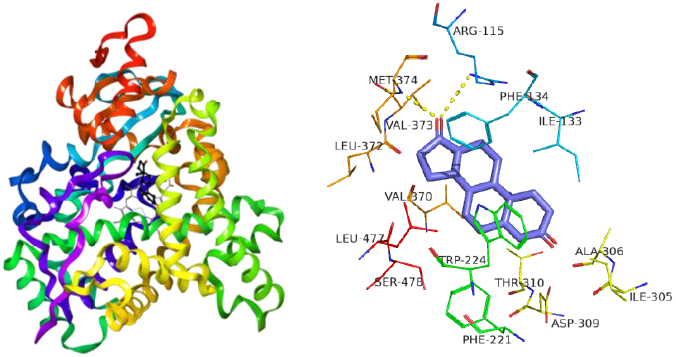
Crystal Structure of Aromatase (CYP19A1 & Protein-ligand interaction ofCYP19Al and ASD

**Figure 2:**
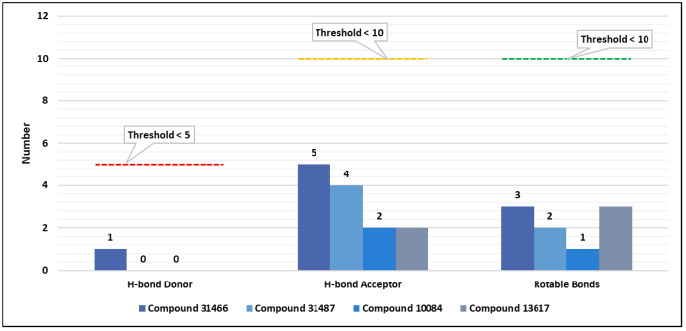
ADME/Tox evaluation

**Figure 3:**
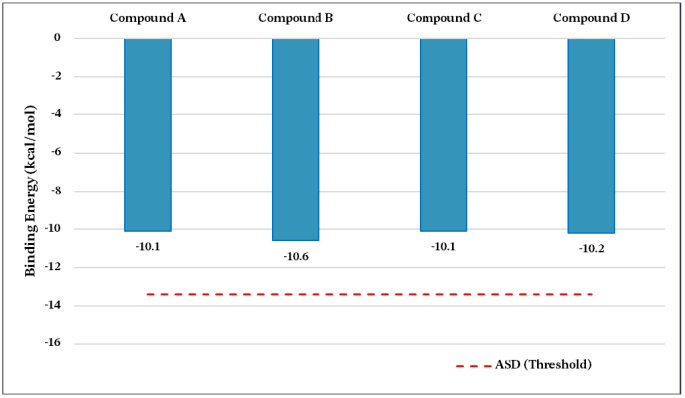
Selection of top hits based on binding affinity with CYP19Al

**Figure 4:**
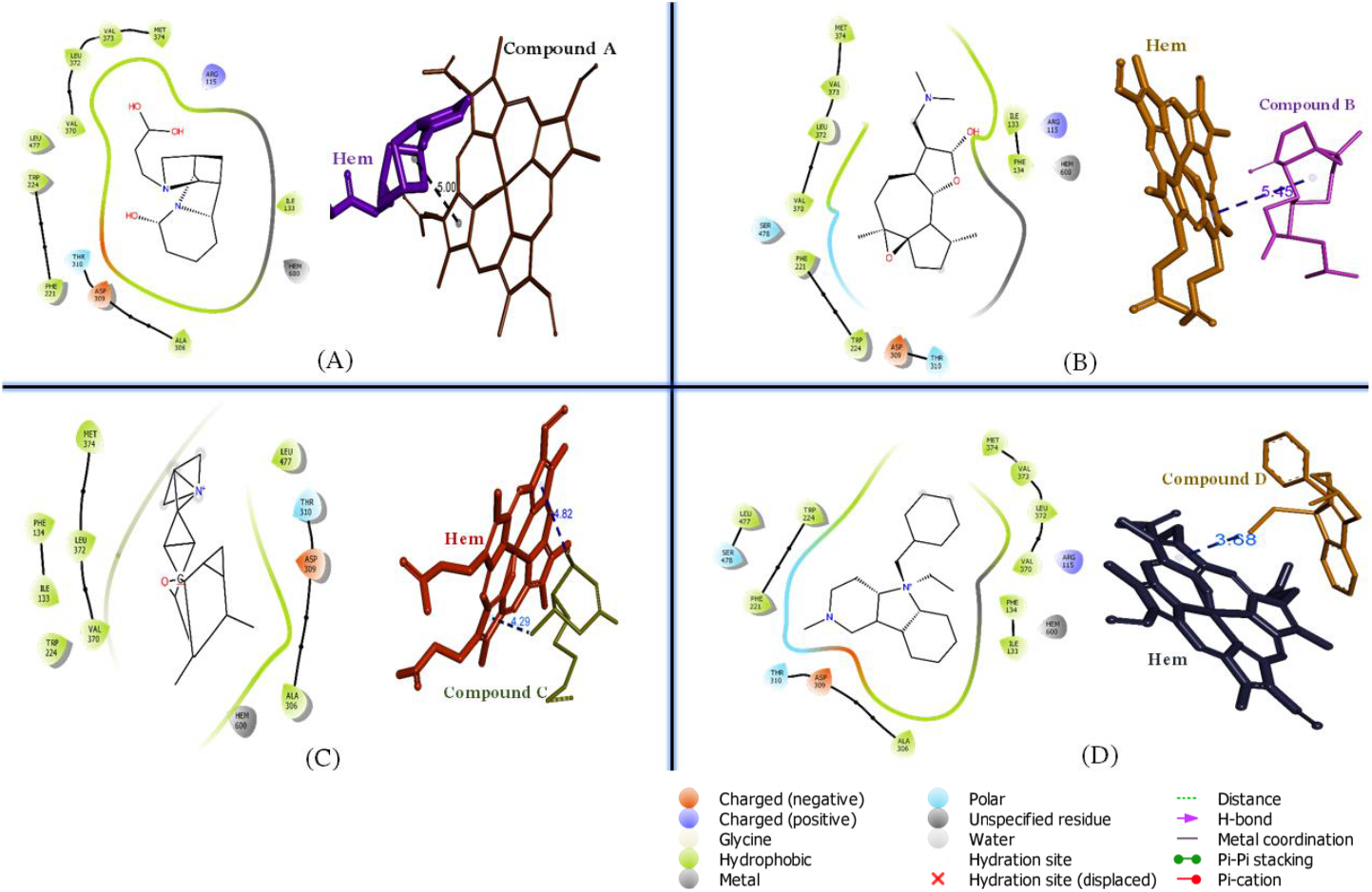
(A) CYP19A1 residues involved in compound A and the compound interaction with hem, (B) CYP19A1 residues involved in compound B and the compound interaction with hem, (C) CYP19A1 residues involved in compound C and the compound interaction with hem, (D) CYP19A1 residues involved in compound D and the compound interaction with hem

### Ligand Profiling

Although chemical synthesis libraries have established chemical functions for proteins, considering kinase inhibitors (ATP analogs) and indole scaffolds, their filtering is challenging. Moreover, competitive inhibition results from a molecule of a similar kind if it is involved in steroid metabolism. Natural Products (NPs) are now one of the most crucial sources in modern pharmacology for creating novel lead compounds and scaffolds. For instance, between 1981 and 2014, more than 50% of newly developed medications were derived from NPs (Newman and Cragg, 2016). After undergoing ADMET filtering, out of the initial 2,24,205 natural compounds retrieved, only 35,234 ligands remained suitable for further docking simulations after their energy was minimized.

### Virtual Screening and ADME Analysis

Androstenedione (ASD) exhibited a binding affinity of -13.4 Kcal/mol. A stronger ligand binding to the target corresponds to a lower negative value of binding affinity (Dallakyan and Olson, 2015). Only 53 out of 9,977 tested ligands had binding energies equal to ASD, indicating they would not cause hormonal imbalance by inhibiting aromatase. A list of these ligands is included in the supplementary data (*S1 Table)*. This suggests that both the ligand and ASD would interact cohesively, maintaining the expression of aromatase at the threshold level.

The four top hits were chosen based on their compliance with ADME/Tox assessment and their proximity to ASD. The compounds labeled A, B, C, and D are 31466, 31487, 10084, and 13617, respectively. Compound B showed a close affinity for the ASD ligand of CYP19A1 and passed the ADME/Tox evaluation.

### Protein-Ligand Interaction Analysis

The top hits must have binding affinities comparable to those of the reference ligand ASD because of the participation of similar amino acid residues and potential binding interactions of various kinds. HEM and the ligand could both accommodate in the binding site, with the ligand improving stability by associating with HEM instead of the protein. Most compounds formed the Pi-alkyl bond type from HEM’s pyrrole ring. The supplementary data include details on the interactions of CYP19A1 and HEM *(SI Figure)* and the nature of the chemical interaction between top hits and CYP19A1 (S2 *Table)*.

Compound B was found to have a higher binding affinity, potentially due to its higher number of hydrophobic residues compared to compounds A and C whose supporting evidence is provided in supplementary data (S2 *Figure)*. Despite having fewer hydrophobic interacting residues than compound D, compound B was selected for its strong affinity for the ASD of CYP19A1 and its passing of the ADME/Tox test.

### Liver Toxicity & Metabolism Assessment

S-adenosylmethionine (SAM), the main biological methyl donor, is synthesized by the liver-specific enzyme, methionine adenosyltransferase (MAT) 1A (Takahashi & Fukusato, 2017). SAM is typically used in transmethylation processes where its methyl group is transferred to various biological acceptors, leading to the creation of S-adenosyl homocysteine (SAH) (Finkelstein, 1990). The binding energy (B.E.) of SAM & SAH was found to be -7.2 kcal/mol. Compound A was not further considered the top hit following its inhibition of human MAT1A.

Cytochrome P450 enzymes function to catalyze a wide range of reactions, many of which are critically important for drug response (Guengerich, 2021). The inability of the top hits to inhibit the Phase I drug-metabolizing enzymes, while contrasting to the binding energy of its respective reference inhibitor, assured to not hinder drug metabolism and all other body functions for which these enzymes are responsible, allowing for the clearance of drugs along with our compound once it enters the body *(Figure 6)*. SwissADME server provided further evidence *(Table 2)*.

**Figure 5:**
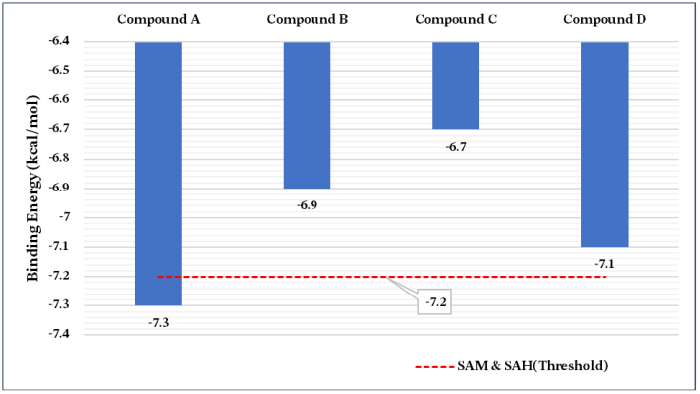
Liver toxicity assessment through SAM & SAH inhibitors binding energy

**Figure 6:**
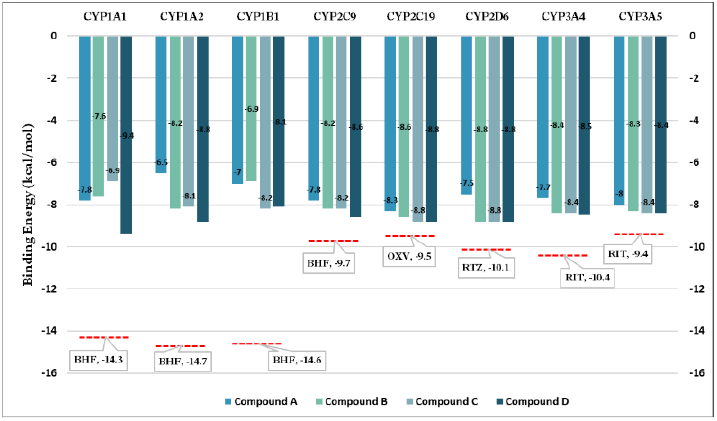
Selection of the best drug candidate based on binding affinity with Phase-I drug-metabolizing enzymes [BHF, OXV, RTZ & RIT are the native/reference ligands of the enzymes CYP1A1, CYP1A2, CYP1B1, CYP2C9, CYP2C19, CYP2D6, CYP3A4, CYP3A5 respectively.]

**Table 1:** Structure validation for each protein using the SAVES v6.0 and ProSA web server.

| Target Protein | Retrieval Source | Z-score | Ramachandran residues | Errat (overall quality factor) |
| --- | --- | --- | --- | --- |
| Aromatase (CYP19A1) | RCSB PDB | -9.24 | 86.6% residues in most favored regions | 97.0721 |
|  |  |  | 11.6% residues in additional allowed regions |  |
|  |  |  | 1.5% residues in generously allowed regions |  |
|  |  |  | 0.2% residues in disallowed regions |  |

**Table 2:** Pharmacokinetics of Compound B via SwissADME.

| Parameters | Pharmacokinetics |
| --- | --- |
| GI absorption | High |
| CYP1A2 inhibitor | No |
| CYP2C19 inhibitor | No |
| CYP2C9 inhibitor | No |
| CYP2D6 inhibitor | No |
| CYP3A4 inhibitor | No |

**Table 3:**
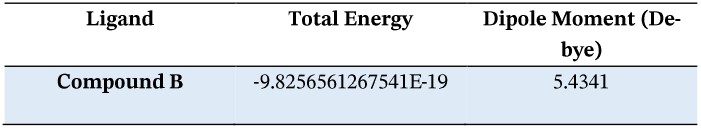
Calculated parameters of the best drug candidate.

### Electronic Structure Study

A molecule’s higher overall energy indicates that it is less stable and more reactive (Rahman et al. 2020). Additionally, free energy is a crucial parameter for describing the interactions of binding partners, where both the sign and magnitude are crucial for expressing the likelihood of bimolecular events, and higher negative values denote superior thermodynamic features (Tripathi et al., 2022).

When compared to drug-like compounds, which have a dipole moment of less than 10 Debye (Pereira & Aires-De-Sousa, 2018), the dipole moment for the chosen compound B was 5.4341 Debye. Furthermore, the molecule exhibited a total energy of -9.8256561267541E-19, indicating its stable state and no requirement for extra energy to support spontaneous binding.

The Molecular Electrostatic Potential (MEP) is directly related to the electronic density of a molecule, and it provides valuable insights into potential sites for electrophilic attack and nucleophilic reactions, as well as hydrogen bonding interactions (Scrocco & Tomasi, 1978) (Luque et al., 2000). The MEP surface shows that the red area has a partially negative charge and is rich in electrons. On the other hand, the blue area lacks electrons and has a partial positive charge. The yellow zone is somewhat electron-rich, while the light blue region is slightly electron-deficient. Finally, the green area is neutral.

As observed in *Figure 7*, the region exhibiting a negative electrostatic potential is located near the oxygen atom of the dioxatetracyclo region, whereas the regions that present a positive potential are localized near the methyl groups of the dioxatetracyclo region.

**Figure 7:**
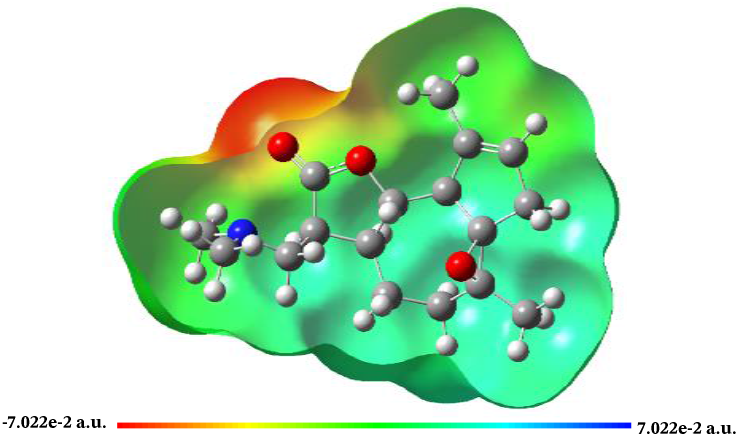
Molecular electrostatic potential map of compound B

A molecule with a small frontier orbital gap is generally associated with high chemical reactivity and low kinetic stability and is also termed a soft molecule (Fukui, 1982) (Gunasekaran et al., 2008). The electronic excitation energy necessary to determine the molecular reactivity and stability of compounds is indicated by the HOMO-LUMO gap energy, which is the difference between the HOMO and LUMO energies (Zhan et al., 2003). The ability of electrons to migrate from the HOMO to the LUMO is negatively impacted by large energy gaps, which ultimately results in poor affinity of the ligand for the target protein (Ali et al., 2022).

The Frontier Molecular Orbitals HOMO and LUMO of compound B in *Figure 8* indicate that it has a low energy gap, making it a soft molecule. Additionally, it has a high HOMO energy (-0.19777 eV), making it the best electron donor. Therefore, compound B was determined to be reactive. The smaller frontier orbital gap of the compound makes it more polarizable and is associated with its high chemical reactivity, low kinetic stability, and softness.

**Figure 8:**
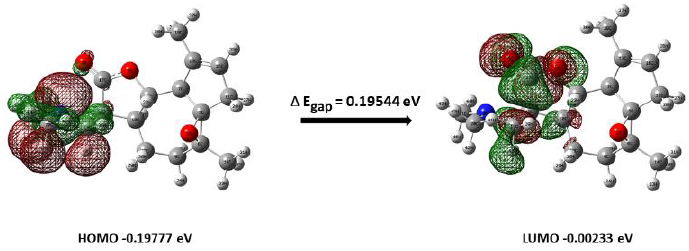
Energy differences (LUMO-HOMO) of the compound B

## Conclusion

Amidst the widespread prevalence of menorrhagia among women, the imperative for drugs devoid of side effects in reproductive health has become evident. In this context, the role of aromatase, particularly its overexpression in heavy uterine bleeding, has emerged as a plausible target of interest. With the advent of computational power, the adoption of computer-aided drug design for prescreening compounds holds the promise of mitigating the likelihood of experimental failures.

Based on the findings of this study, a computational approach can be used to efficiently identify potential therapeutic candidates for regulating hormone levels. This research identified a highly promising molecule, (lR,3S,6S,7S,10S,HR)-7-[(dimethylamino)methyl]-3,12-di-methyl-2,9-dioxatetracyclo [9.3.0.01,3.06,10] tetradec-12-en-8-one, which possesses the ability to maintain aromatase, a critical enzyme in steroidogenesis, at the desired level. Moreover, the compound boasts a favorable ADMET profile and does not impede SAM production, further demonstrating its potential to serve as a medicinal treatment. Armed with this knowledge, it may be possible to rapidly identify drug-like molecules, expedite their use in clinical trials, and assess their effectiveness in treating female hormonal disorders.

Through research and efforts to tackle the identified issues, we aim to create a safer and more effective treatment option for women with menorrhagia, ultimately leading to improved healthcare and quality of life for many patients.

## Materials and Methods

### Protein and Ligand Preparation

The crystallographic structure with 5 A resolution was obtained in the PDB format from the RCSB Protein Data Bank (https://www.rcsb.org/) which was analyzed for its z-score online server proSA (https://prosa.ser-vices.came.sbg.ac.at/prosa.php) and the Ramachandran plot analysis was done using another server SAVES v6 (https://saves.mbi.ucla.edu/). A set of amino acid residues were chosen as a ligand binding site for molecular docking simulations with the aid of PyMOL. Before docking, the protein underwent some preparations, including the addition of hydrogen atoms, the combination of non-polar bonds, the inclusion of Gasteiger charges, and the conversion of the protein file format into PDBQT using AutoDock tools (Morris et al., 2009).

It was rational to examine the natural product library of the Zincl5 database (https://zincl5.docking.org/substances/home/). The OSIRIS program was used to predict the toxicity profiles and drug-likeness of potential lead compounds (Rashid, 2020). The fundamental element crucial for rational drug design is the implementation of Lipinski’s Rule of 5 (RO5) concerning bioactive compounds. This rule was employed using the Osiris Data Warrior program to assess the suitability of the established compounds. Energy minimization was performed with a universal force field (UFF) using the conjugate gradient algorithm in OpenBabel GUI, which is available at the PyRx interface (Dallakyan, 2008). Ligand preparation was carried out in Openbabel GUI (O’Boyle et al., 2011) using PyRx 0.9.8 setup by energy minimization using a force field and converted to PDBQT file format to be used for docking.

### Molecular Docking and Protein-Ligand Interaction Analysis

Structure-based virtual screening was conducted with the primary goal of avoiding partiality among classes of molecules. The Aromatase enzyme (PDB: 3EQM) was docked with the 35,234 compounds obtained from the ADMET filtration and energy minimization process. Docking was performed in the virtual screening software PyRx 0.9.8 platform (https://pyrx.sourceforge.io/) using AutoDock Vina (Trott & Olson, 2009). The top hits were identified based on their proximity to CYP19A1 negative binding energy and druglike properties (number of H-bond donors and acceptors, rotatable bonds) analysis.

The amino acid residues involved in making possible interactions with the protein structure, bond length, types of interaction with the top hit ligands were observed through PyMOL (Seeliger & De Groot, 2010), Discovery Studio Visualizer (BIOVIA, Dassault Systemes, 2021), Ligplot+ (LigPlot+ Version 2.2) and Schrodinger Maestro (Schrodinger Maestro, 4.6.0) software.

### Toxicity and Clearance Analysis

The top hits were docked with the hMATlA protein, where SAM (S-Adenosylmethionine) was used as a reference inhibitor. Ligands with a B.E. greater than that of the SAM inhibitor, which could replace the SAM of HMATlA, were filtered off and removed from the ligand library prepared for molecular docking (Ghimire et al., 2022).

Additionally, the top hits were subjected to another screening with Phase-I drug-metabolizing enzymes to ensure their successful clearance from the body, where the respective reference or native ligands were used as reference inhibitors for screening. Ligands with a B.E. greater than that of the reference inhibitors, which could replace the inhibitors, were not considered top hits. This was further cross checked in SwissADME server (http://swissadme.ch/) of Swiss Institute of Bioinformatics.

### Density Function Theory (DFT)

DFT calculations were performed using the GAUSSIAN 03 platform to interpret the atomic arrangement of the studied compound (Mohapatra et al., 2021). It was performed using functional B3LYP with the 6-31G** basic set in Gaussian 03 (Zandler & D’Souza, 2006) to analyze the Dipole Moment, Molecular Electrostatic Potential (MEP) and Frontier Molecular Orbitals (FMO).

The molecular electrostatic potential of the title compound was analyzed to identify its reactive sites using. The analysis of Frontier Molecular Orbitals (FMO) describes one-electron excitation from the highest occupied molecular orbital (HOMO) to the lowest unoccupied molecular orbital (LUMO). The HOMO energy is directly related to the ionization potential, and the LUMO energy is related to the electron affinity. The HOMO and LUMO energy gaps explain the eventual charge transfer interactions within the molecules.

## Acknowledgment

We gratefully acknowledge the generous support from the Tribhuvan University Biotechnology Department, including access to their computational facilities and valuable insights from the alumni students of Professor Dr. Pramod Aryal.

Thanks to Prof. Dr. Rajendra Prasad Koirala for encouraging us to further expand the scope of our study to include MD simulation. Our sincere thanks are also extended to Prof. Dr. Rameshwor Adhikari and RECAST for providing us with a conducive and peaceful environment for conducting our preliminary literature mining, which was vital to the success of this project.

