## Supplemental Table 1 for "Computational Assessment of Aromatase: Unmasking Therapeutic Avenues for Female Hormonal Disorders"

**Virtual Screening & Top Hits Identification**

| **Molecule** | **MW** | **cLogP** | **cLogS** | **H-a** | **H-d** | **TPSA** | **Drug-**  **likeness** | **RB** | **B.E. with ASD (Kcal/mol)** |
| --- | --- | --- | --- | --- | --- | --- | --- | --- | --- |
| Compound 10041 | 360.416 | 2.2396 | -3.839 | 6 | 2 | 77.56 | 4.7077 | 6 | -10.3 |
| **Compound 10084** | **258.384** | **-1.2295** | **-2.115** | **2** | **0** | **13.11** | **0.083412** | **1** | **-10.1** |
| Compound 10217 | 302.373 | 2.1979 | -2.637 | 5 | 1 | 58.64 | 7.2369 | 3 | -10.0 |
| Compound 10658 | 264.283 | 1.3523 | -3.273 | 4 | 3 | 65.12 | 3.0996 | 1 | -10.4 |
| Compound 11478 | 286.37 | 3.8064 | -3.958 | 3 | 0 | 27.69 | 0.1613 | 1 | -10.9 |
| **Compound 13617** | **305.444** | **2.5733** | **-3.326** | **2** | **0** | **3.24** | **4.5723** | **3** | **-10.2** |
| Compound 13738 | 330.422 | 2.8453 | -3.779 | 4 | 1 | 59.67 | 0.45912 | 3 | -10.4 |
| Compound 14251 | 428.491 | 2.2844 | -2.949 | 7 | 1 | 74.65 | 3.9312 | 7 | -10.6 |
| Compound 14868 | 281.351 | 0.8704 | -2.9 | 5 | 1 | 86.46 | 0.82741 | 3 | -10.1 |
| Compound 15070 | 372.427 | 2.0842 | -3.383 | 6 | 1 | 73.07 | 2.5087 | 6 | -10.4 |
| Compound 15159 | 275.347 | 1.5705 | -3.05 | 4 | 3 | 69.56 | 2.8523 | 3 | -10.0 |
| Compound 1746 | 266.336 | 1.0996 | -2.213 | 4 | 2 | 66.76 | 1.1448 | 0 | -10.4 |
| Compound 18103 | 262.304 | 1.6067 | -2.351 | 4 | 1 | 63.6 | 0.39471 | 0 | -10.1 |
| Compound 19871 | 276.335 | 1.0381 | -1.752 | 5 | 2 | 53.96 | 1.3847 | 2 | -10.2 |
| Compound 20159 | 275.303 | 1.1202 | -3.886 | 5 | 2 | 70 | 0.22969 | 2 | -10.0 |
| Compound 2285 | 278.259 | -0.5448 | -2.064 | 6 | 1 | 82.06 | 1.6836 | 2 | -10.3 |
| Compound 2482 | 287.314 | 1.2078 | -2.776 | 5 | 2 | 62.16 | 3.751 | 0 | -10.4 |
| Compound 2634 | 342.39 | 2.9165 | -3.303 | 5 | 1 | 64.99 | 0.39139 | 6 | -10.2 |
| Compound 3076 | 264.327 | 2.3287 | -2.672 | 3 | 1 | 48.02 | 3.0807 | 4 | -10.2 |
| Compound 3282 | 272.255 | 1.8017 | -3.103 | 5 | 3 | 86.99 | 0.29065 | 0 | -10.3 |
| Compound 3373 | 297.353 | 1.678 | -2.274 | 4 | 2 | 60.77 | 3.6726 | 3 | -10.0 |
| Compound 5432 | 258.36 | 3.3493 | -2.951 | 2 | 1 | 29.46 | 0.20749 | 1 | -10.1 |
| Compound 6482 | 240.305 | 1.8465 | -2.826 | 3 | 1 | 36.1 | 5.8984 | 1 | -10.0 |
| Compound 6683 | 307.392 | 3.3192 | -3.733 | 3 | 1 | 38.33 | 1.4246 | 4 | -10.0 |
| Compound 7261 | 306.448 | 2.2455 | -1.719 | 4 | 0 | 32.78 | 2.0682 | 2 | -10.4 |
| Compound 7576 | 314.344 | -0.0495 | -2.575 | 7 | 0 | 81.73 | 1.3169 | 2 | -10.3 |
| Compound 7990 | 273.375 | 2.3934 | -2.507 | 3 | 0 | 31.35 | 0.87945 | 1 | -10.2 |
| Compound 8235 | 272.391 | 1.9955 | -1.708 | 3 | 1 | 26.71 | 4.1277 | 1 | -10.4 |
| Compound 8298 | 298.428 | 3.048 | -2.299 | 3 | 0 | 23.55 | 3.9051 | 3 | -10.0 |
| Compound 8340 | 300.401 | 2.3584 | -1.659 | 4 | 1 | 43.78 | 3.8865 | 3 | -10.3 |
| Compound 8345 | 286.417 | 2.5604 | -1.904 | 3 | 1 | 26.71 | 3.7885 | 3 | -10.0 |
| Compound 8373 | 272.347 | 1.4496 | -1.119 | 4 | 1 | 43.78 | 3.822 | 1 | -10.2 |
| Compound 8376 | 256.348 | 1.7953 | -1.415 | 3 | 0 | 23.55 | 3.8345 | 1 | -10.2 |
| Compound 8491 | 351.492 | 3.2595 | -3.859 | 4 | 2 | 44.37 | 6.1454 | 6 | -10.1 |
| Compound 28814 | 250.337 | 2.2975 | -2.194 | 3 | 2 | 57.53 | 0.81055 | 0 | -10.2 |
| Compound 30337 | 250.337 | 1.9449 | -2.813 | 3 | 1 | 46.53 | 1.9981 | 0 | -10.2 |
| Compound 31156 | 251.372 | 4.1807 | -3.695 | 1 | 1 | 12.03 | 2.1478 | 3 | -10.6 |
| **Compound 31466** | **262.308** | **-0.7049** | **-0.987** | **5** | **1** | **60.85** | **3.4282** | **3** | **-10.1** |
| **Compound 31487** | **291.39** | **0.7985** | **-2.003** | **4** | **0** | **42.07** | **3.067** | **2** | **-10.6** |
| Compound 31598 | 251.372 | 4.1807 | -3.695 | 1 | 1 | 12.03 | 2.1478 | 3 | -10.4 |
| Compound 31612 | 307.352 | 2.0903 | -3.313 | 5 | 2 | 67.01 | 4.3896 | 5 | -10.2 |
| Compound 31745 | 301.301 | 2.0443 | -2.519 | 7 | 0 | 77.69 | 5.1095 | 3 | -10.5 |
| Compound 31747 | 337.465 | 3.0037 | -3.357 | 4 | 2 | 44.37 | 3.9468 | 7 | -10.4 |
| Compound 31934 | 288.386 | 3.2793 | -2.969 | 3 | 1 | 38.69 | 0.35982 | 2 | -10.3 |
| Compound 32263 | 285.298 | 0.7328 | -2.637 | 5 | 3 | 78.79 | 2.9918 | 2 | -10.6 |
| Compound 32264 | 285.298 | 0.7328 | -2.637 | 5 | 3 | 78.79 | 2.9918 | 2 | -10.7 |
| Compound 34625 | 306.708 | 1.3659 | -3.573 | 7 | 0 | 79.13 | 1.3587 | 2 | -10.9 |
| Compound 34777 | 359.396 | 2.2536 | -3.62 | 5 | 3 | 78.79 | 1.3922 | 6 | -10.6 |
| Compound 35651 | 255.272 | 0.8028 | -2.619 | 4 | 3 | 69.56 | 2.9261 | 1 | -10.4 |
| Compound 35665 | 326.347 | 2.3552 | -3.581 | 5 | 3 | 94.83 | 1.6525 | 7 | -10.1 |
| Compound 35675 | 299.369 | 2.8892 | -3.785 | 4 | 0 | 38.77 | 1.6316 | 0 | -10.1 |
| Compound 109 | 307.476 | 2.2071 | -3.951 | 3 | 3 | 66.48 | 0.7098 | 0 | -10.1 |
| Compound 564 | 409.484 | 1.781 | -2.687 | 7 | 1 | 78.95 | 2.2375 | 7 | -10.1 |

Table S1: Ligands with Binding Energy (B.E) close to the natural ligand ASD for CYP19A1 and their respective ADMET properties
