## Supplemental Table 2 for "Computational Assessment of Aromatase: Unmasking Therapeutic Avenues for Female Hormonal Disorders"

**Protein-Ligand Interaction**

| Ligands | Bond Length | Bond Type | Chemical Group of Hem |
| --- | --- | --- | --- |
| Compound 31466 | 5.0 Å | Pi – alkyl | Pyrrole ring |
| Compound 31487 | 5.34 Å | Pi – alkyl | Pyrrole ring |
| Compound 10084 | 4.82 Å | Pi – alkyl | Pyrrole ring |
|  | 4.29 Å | Pi – alkyl | Pyrrole ring |
| Compound 13617 | 3.68 Å | Pi – sigma | Pyrrole ring |
| ASD (native) | 4.93 Å | Pi – alkyl | Pyrrole ring |
|  | 4.02 Å | Pi – alkyl | Pyrrole ring |
|  | 4.26 Å | Alkyl | Methyl group |

S2 Table 1: Nature of chemical interaction between top hits and CYP19A1
