## Supplemental Figure 1 for "Computational Assessment of Aromatase: Unmasking Therapeutic Avenues for Female Hormonal Disorders"

**Protein and HEM Interaction**

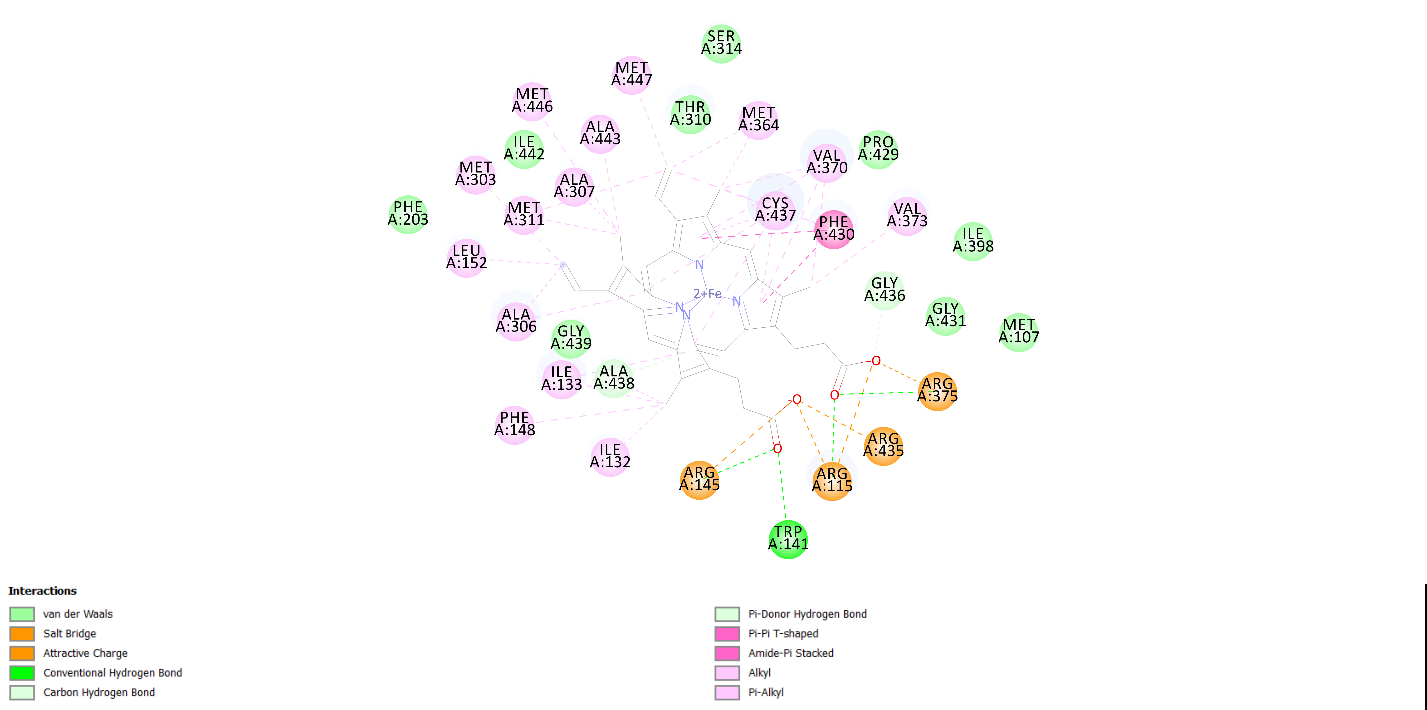

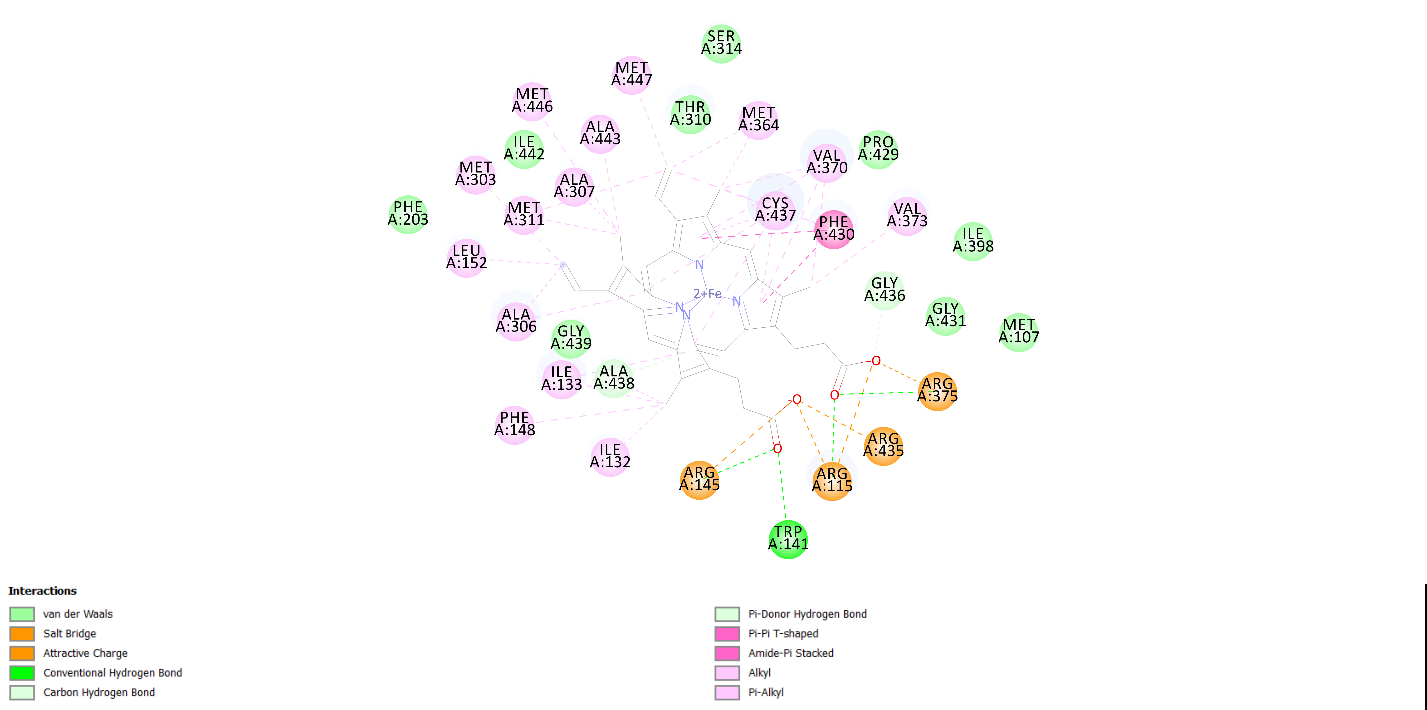

S1 Figure 1: 2D interaction between Hem and CYP19A1 as visualized in BIOVIA Discovery Studio Visualizer
