## Supplemental Figure 2 for "Computational Assessment of Aromatase: Unmasking Therapeutic Avenues for Female Hormonal Disorders"

**Hydrophobic Interactions**

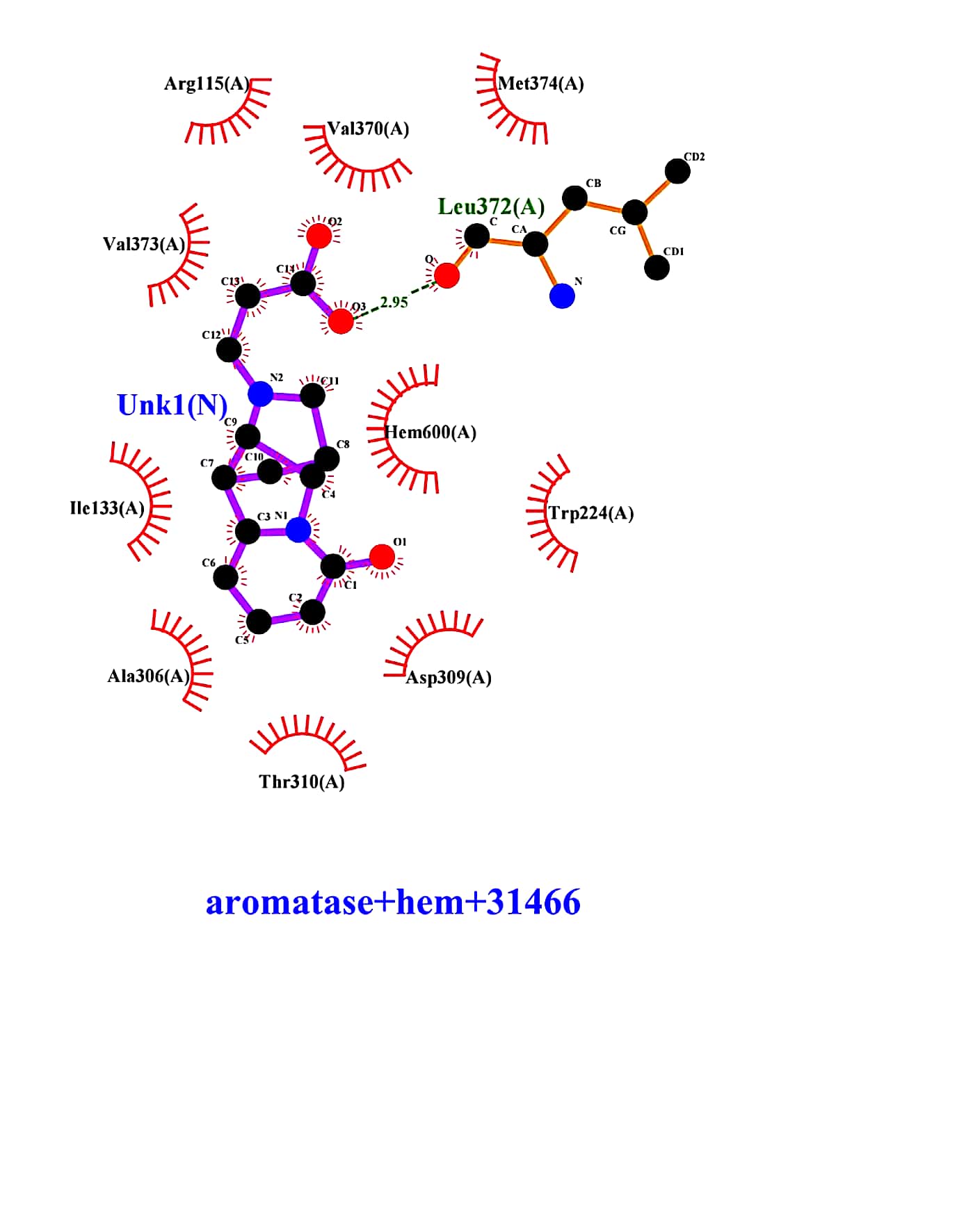

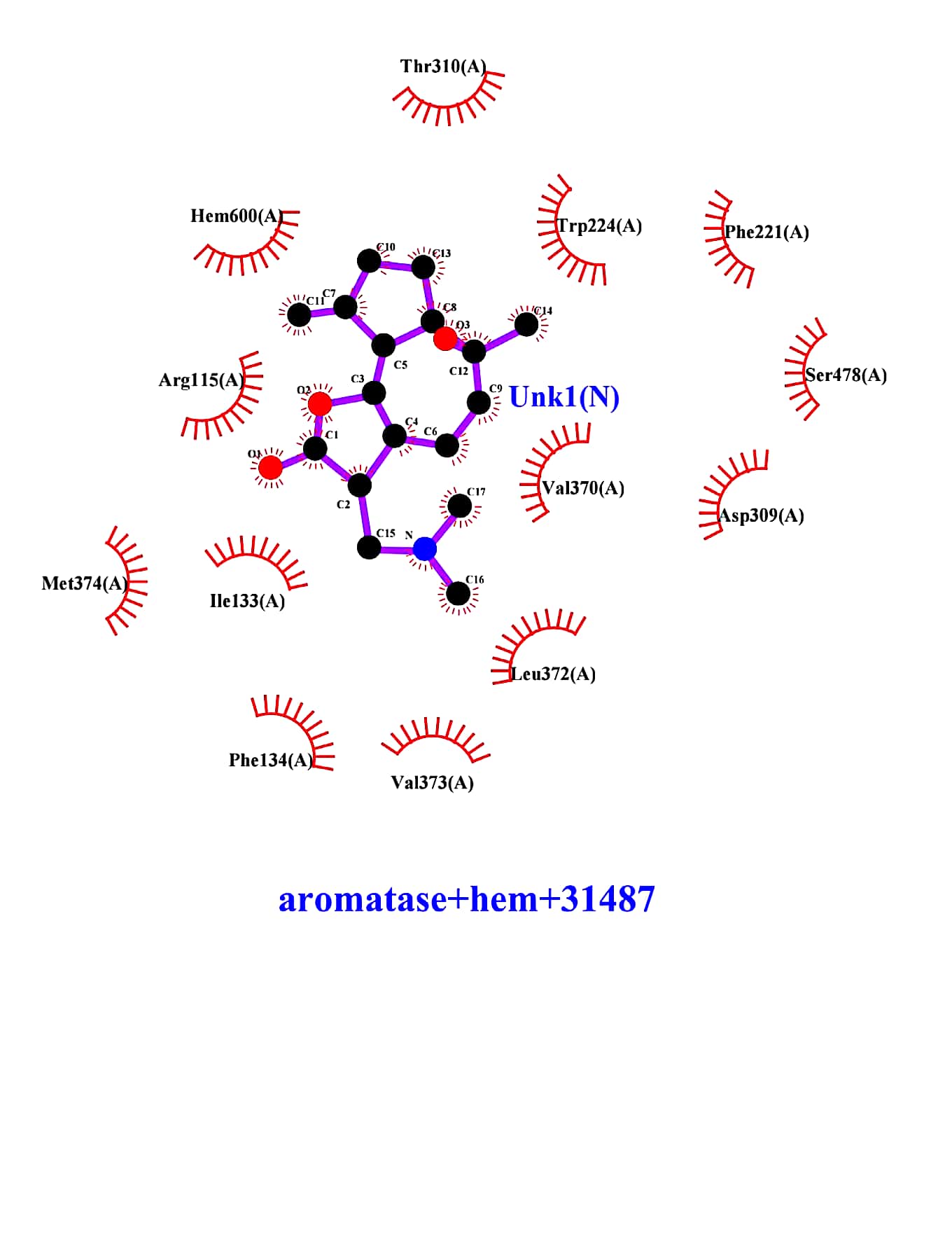

S2 Figure 2: Hydrophobic Interactions of Compound 31487 with CYP19A1

S2 Figure 1: Hydrophobic Interactions of
Compound 31466 with CYP19A1

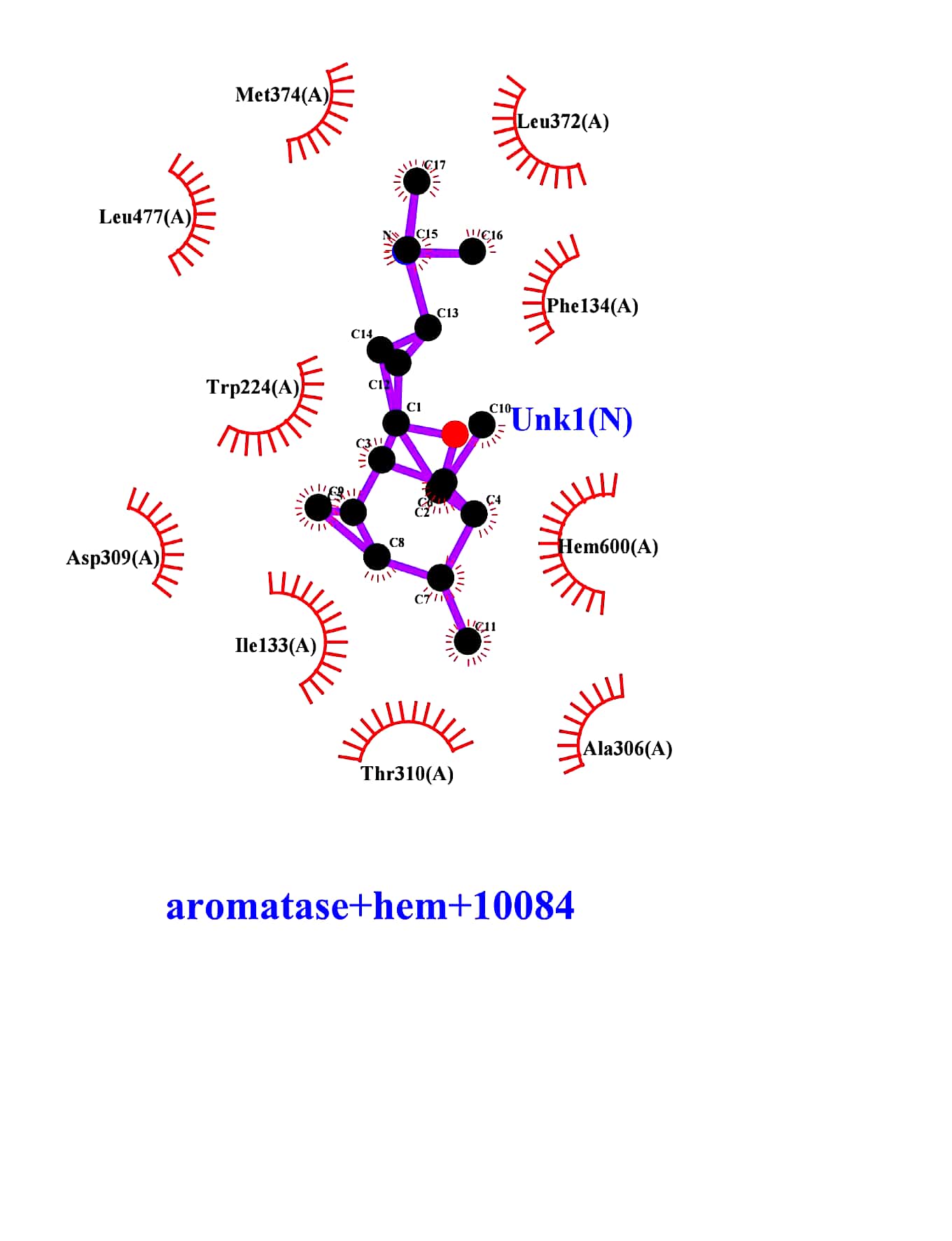

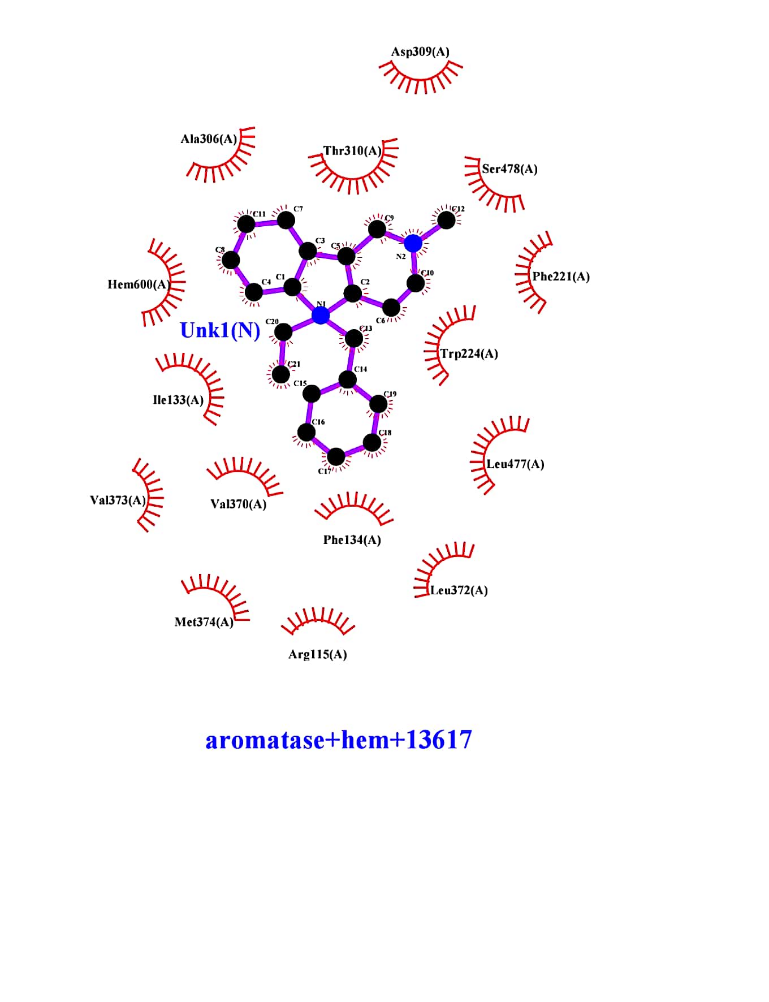

S2 Figure 3: Hydrophobic Interactions of Compound 10084 with
CYP19A1

S2 Figure 4: Hydrophobic Interactions of Compound 13617 with CYP19A1
